# UbE3-APA: A Bioinformatic Strategy to Elucidate Ubiquitin E3 Ligase Activities in Quantitative Proteomics Study

**DOI:** 10.1101/2022.01.16.476541

**Authors:** Yao Gong, Yue Chen

## Abstract

**Motivation:** Ubiquitination is widely involved in protein homeostasis and cell signaling. Ubiquitin E3 ligases are critical regulators of ubiquitination that recognize and recruit specific ubiquitination targets for the final rate-limiting step of ubiquitin transfer reactions. Understanding the ubiquitin E3 ligase activities will provide knowledge in the upstream regulator of the ubiquitination pathway and reveal potential mechanisms in biological processes and disease progression. Recent advances in mass spec-trometry-based proteomics have enabled deep profiling of ubiquitylome in a quantitative manner. Yet, functional analysis of ubiquitylome dynamics and pathway activity remains challenging.

**Results:** Here, we developed a UbE3-APA, a computational algorithm and stand-alone python-based software for Ub E3 ligase Activity Profiling Analysis. Combining an integrated annotation database with statistical analysis, UbE3-APA identifies significantly activated or suppressed E3 ligases based on quantitative ubiquitylome proteomics datasets. Benchmarking the software with published quantitative ubiquitylome analysis confirms the genetic manipulation of SPOP enzyme activity through overexpression and mutation. Application of the algorithm in the re-analysis of a large cohort of ubiquitination proteomics study revealed the activation of PARKIN and the co-activation of other E3 ligases in mitochondria depolarization-induced mitophagy process. We further demonstrated the application of the algorithm in the DIA-based quantitative ubiquitylome analysis.

**Availability:** Source code and binaries are freely available for download at URL: https://github.com/Chenlab-UMN/Ub-E3-ligase-Activity-Profiling-Analysis, implemented in python and supported on Linux and MS Windows

**Supplementary information:** Supplementary data are available.

## 1 Introduction

Ubiquitylation is a key protein post-translational modification (PTM) involved in diverse cellular processes including protein homeostasis, cell signaling and epigenetic regulations. Its E1-E2-E3 cascades linking an iso-peptide bond between the c-terminus of ubiquitin and a lysine residue of the target protein (Pickart, 2003) to form a mono- or a polymer chain of ubiquitin with eight distinct linkage types. Ubiquitylation not only acts as the essential modification in protein degradation through proteasome that accounted for the breakdown of over 80% of the proteins (Lee and Goldberg, 1998), but it also plays a crucial role in non-degradative functions, including regulation of protein trans-location, protein-protein interactions and enzymatic activity (Schnell and Hicke, 2003). Within the process of ubiquitylation, E3, also known as the ubiquitin ligase, mediates the ubiquitination substrates specificity (Ordureau *et al.*, 2015; Pickart, 2003). Changes in the E3 ligase activities will lead to changes in ubiquitination of its target proteins, and further regulate various downstream cellular processes including cell-cycle, apoptosis, and transcription regulation (Hoeller and Dikic, 2009). Studies have found that dysfunction of E3 ligases and these cellular functions may lead to neurodegenerative diseases (McNaught *et al.*, 2001; Oddo, 2008), cardiovascular diseases (Herrmann *et al.*, 2004), and development of cancer (Hoeller *et al.*, 2006; Nakayama and Nakayama, 2006), while therapies and drugs have been developed to target specific E3 ligases for potential clinical applications (Petroski, 2008; Bulatov and Ciulli, 2015). Therefore, it is important to develop strategies that evaluate the activities of different E3 ligases in a system-wide manner.

Recent advances in mass spectrometry have enabled deep profiling of PTM pathways. Combining with quantitative proteomics strategies such as SILAC and isobaric labeling, proteomics analysis allows system-wide profiling of PTM dynamics at the site-specific level. Such quantitative information provides a rich resource to develop computational tools evaluating PTM pathway activities (Olsen and Mann, 2013). Recent efforts studying kinase activities based on quantitative phosphorylation datasets have led to the development of several tools, including PTMsigDB (Krug *et al.*, 2019), IKAP (Mischnik *et al.*, 2016), KinasePA (Yang *et al.*, 2016), KSEA (Wiredja *et al.*, 2017), and KEA3 (Kuleshov *et al.*, 2021). Among these models, KEA3 collected 24 kinase substrate libraries from different sources as their database and test significance of kinase integrating sum rank tests results of all libraries, while the rank-sum test in PTMsigDB was supported by a collection of site-specific PTM signature of perturbations, kinase states and pathway activities from published studies (Krug *et al.*, 2019). IKAP used a non-linear optimization routine to find enriched kinase, KinasePA used the direction pathway analysis to study insulin pathways (Yang *et al.*, 2014) and KSEA applies z-score test to find differentially activated kinases. Despite these advances in kinase analysis, there is lack of bioinformatic strategies for evaluating ubiquitin E3 ligase activities.

Improvements in biochemical enrichment and chemical labeling strategies have allowed global quantification of ubiquitination dynamics (Kim *et al.*, 2011; Wagner *et al.*, 2011; Udeshi *et al.*, 2013; Elia *et al.*, 2015) and measurement of site-specific ubiquitination stoichiometries (Li *et al.*, 2019). Recent bioinformatic efforts have led to the development of multiple enzyme-substrate databases in the ubiquitination pathway (Li *et al.*, 2017; Nguyen *et al.*, 2016; Li *et al.*, 2021; Chen *et al.*, 2019; Han *et al.*, 2012; Du *et al.*, 2011). In this study, based on an integrated resource of ubiquitin E3 ligase and substrate network, we proposed a computational strategy UbE3-APA, Ubiquitin E3 ligase Activity Profiling Analysis (**Figure 1, Table S1**), for systematic evaluating ubiquitin E3 ligase activity based on quantitative ubiquitylome analysis. The model was validated with two published large-scale proteomics studies with different biological context (Theurillat *et al.*, 2014; Sarraf *et al.*, 2013) and confirmed known regulatory mechanisms in the pathway.

**Figure 1.**
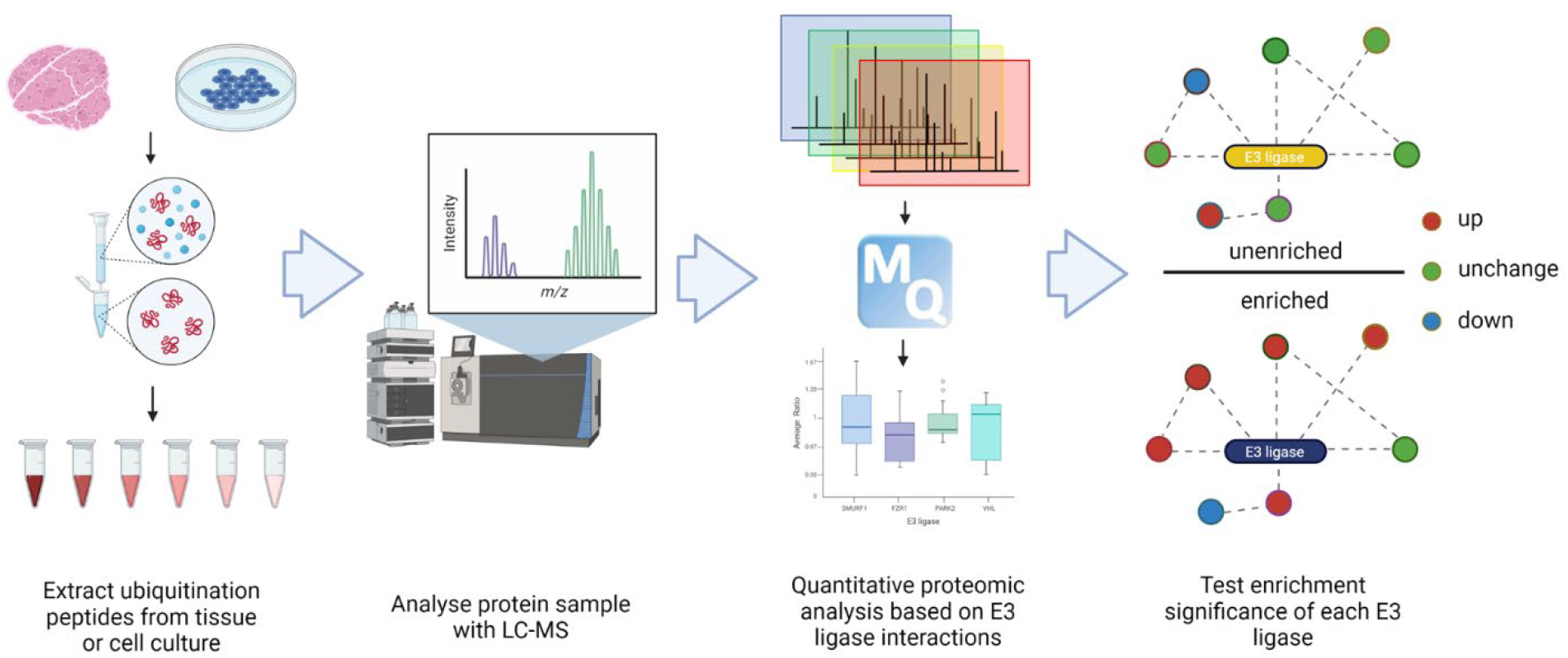
A workflow of analyzing quantitative ubiquitylation proteome with E3 ligases activity profiling analysis.

## 2 Methods

### 2.1 Collecting E3–substrate interactions (ESIs)

We integrated ubiquitin E3 ligases and substrates relationship data from the following three sources: UbiBrowser (Li *et al.*, 2017), Ubinet (Li *et al.*, 2021; Nguyen *et al.*, 2016), and a multidimensional database collection (Chen *et al.*, 2019). UbiBrowser is an extensive database that collects interactions between E3 ligases and substrates. They incorporate ESIs from both literature manual curation of and prediction based on a set of biological features and Bayesian models in their database. Another online platform that updated recently, Ubinet, focuses on ESI collection, prediction, and visualization across different species. They predicted ESIs based on the substrate specificity of E3 ligases extracted from experiment verified interactions. To characterize the interaction network between E3 ligases and their substrates, the Chen group collected ESIs from a variety of sources: E3net (Han *et al.*, 2012), hUbiquitome (Du *et al.*, 2011), Uniprot (Bateman *et al.*, 2017), and BioGRID (Chatr-Aryamontri *et al.*, 2017). They collected ESIs directly from the first two sources, and they gathered the interaction between E3 ligase and proteins in the last two databases through data mining and selected those physical interactions supported by low throughput methods. By integrating the Ub E3-substrate interaction network from these three resources, we established the database for Ub E3 ligase activity profiling analysis (**Table S2**). Only the interactions that were supported by literature from all sources above were integrated into our database. Those interactions solely supported by prediction models were not included.

### 2.2 Algorithm development for ubiquitin E3 ligase activity profiling analysis

The E3 ligase activity analysis model profiles E3 ligase activities based on a bootstrapping procedure by evaluating the difference between the quantitative ratios of E3 ubiquitination targets and the overall background. Firstly, the program collects the quantitative ratios of identified ubiquitination sites and proteins from proteomics analysis. To normalize the site-specific ubiquitination ratios for statistical analysis, the program offers two options – computationally normalized values or protein-normalized values. Computationally normalized ubiquitination ratios are often provided by the quantification software. For example, Maxquant provides normalized site-specific ubiquitination ratios based on the median ratios of all quantified sites. To obtain protein-normalized ubiquitination ratios, the program will fetch the original ratios of both the ubiquitination sites and their corresponding ubiquitination proteins. The protein quantification ratios should be calculated excluding ubiquitinated peptides. The protein-normalized ubiquitination site ratios are calculated by dividing the original site ratios by the original ratios of the corresponding proteins. The normalized site-ratios are then log2 transformed for downstream analysis and averaged to generate the quantification ratios of ubiquitination protein substrates.

Secondly, based on the integrated ubiquitin E3 ligase-substrate database, the program iteratively analyzes each E3 ligase and extract all substrates quantified for each E3 ligase in the quantitative datasets. Then, for each E3 ligase, the program collects the total number of quantified targets and the average quantification ratios of its targets.

Thirdly, the program performs randomized selection from the quantification datasets. At this point, the program offers two options for enrichment testing – protein-level profile analysis and site-level profile analysis. For protein-level profile analysis, the program randomly selects the same number of ubiquitylation proteins as the number of targets for a specific E3 ligase and then computes the average quantification ratios of selected ubiquitination proteins. For site-level profile analysis, the program randomly selects the same number of ubiquitylation sites as the number of sites quantified for known targets of a specific E3 ligase and then computes the average quantification ratios of selected ubiquitination sites. The random selection process is repeated various times for every E3 ligase for parameter optimization and the data analysis in this study was performed with 10,000 repeats.

Lastly, the average ratios of randomly selected groups of ubiquitination proteins or sites were fit into a normal distribution based on the central limit theorem. Based on this distribution, the program calculates the statistical significance for the averaged ratio of the E3 ligase protein targets or sites quantified in the dataset using the formula below.

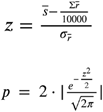

Here 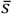 stands for the average quantification ratios of ubiquitination substrate proteins or sites quantified for a given E3 ligase in the dataset, 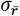 stands for the average ratio of one group of randomly selected proteins or sites, 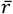 stands for the standard deviation for the distribution of all the average ratios of randomly selected groups of ubiquitination proteins or sites, z stands for z-score and p stands for the p-value.

The program outputs tab-delimited results in text format. To generate a more concise output file, the program offers the option to group E3 ligases. The grouping is helpful as some E3 ligases share ubiquitination targets and depending on the analysis depth, not all targets are quantifiable in the datasets. E3 ligases are grouped together with the leading E3 ligases has all the ubiquitination substrates in this group while the remaining E3 ligases in the group only accounts for a subset of the ubiquitination substrates in the group with no unique substrates. In this way, users can filter out E3 ligases that have no unique substrates but get enriched because of a few common substrates as shown in our analysis result, and therefore, put more emphasis on the E3 ligases whose activity profiles dominate the ubiquitination dynamics. The workflow was written into a python package, and can be accessed from either PyPI, the standard way of installing python package, or our GitHub webpage.

### 2.3 Analysis of large-scale quantitative ubiquitylome proteomics datasets

We benchmarked our algorithm using two published global ubiquitylome analysis. First, we collected original ubiquitylation ratios generated by two studies. One study focused on how SPOP-mutant affects ubiquitylome and prostate cancer (Theurillat *et al.*, 2014). From this study, we collected the protein-normalized median log2 ratio of quantified ubiquitination site under each experimental condition. The other one focused on the relation between mitochondrial depolarization and PARKIN-dependent ubiquitylome (Sarraf *et al.*, 2013). In this study, we collected the log2 site ratios of quantified ubiquitination site under each experimental condition. Second, we reorganized original data into different tables according to experimental groups described in literature. In this way, the SPOP related data was reorganized into six groups, containing two of mutant-control, mutant-wildtype and mutant-wildtype each. Meanwhile, the PARKIN-related data was reorganized into 73 groups of experiments treated with different chemicals or with various genetic background. Thirdly, the protein-level UbE3-APA analysis was applied to profile E3 ligase activities in both studies. Ubiquitin site ratios from both studies were log2 ratios, so we analyzed them directly without further log transformation. For the PARKIN study, the grouped protein-level analysis was performed. In the group mode, the E3 ligases in the results were clustered when they are sharing the same set of quantifiable targets and the E3 ligase that has the greatest number of quantifiable substrates in the group was defined as the leading E3 ligase. Correlation of E3 ligase profiles between each pair of experiments was calculated with the two-dimensional Euclidean distance between E3 ligase activity profiles in experiments.

We further applied our model to two recently published ubiquitylome studies with DIA analysis. First, we gathered original ubiquitylation intensities generated by two studies. One study explored how tumor necrosis factor (TNF) treatment affects ubiquitylome (Hansen *et al.*, 2021a). And we collected the average log2 intensities of quantified ubiquitination site of treated group and mock group from this study respectively. The other study investigated ubiquitylome changes under USP7 inhibition with chemical inhibition and knockdown methods (Steger *et al.*, 2021). In this study, we collected the average log2 site intensities of quantified ubiquitination site under each experimental condition respectively. Second, we reorganized original intensity data into ratios by comparing different treatment groups described in literature. In this way, we calculated the Treated/Mock ratios in TNF treatment study. And we calculated siCTRL+FT671 / siCTRL+DMSO and siUSP7+FT671 / siUSP7+DMSO in the USP7 study. Lastly, we applied protein-level UbE3-APA analysis for both studies in the grouped mode.

### 2.4 Model accessibility and utility

We packed the whole UbE3-APA model into a python3 library on PyPI to make it accessible. And the most direct way of installing the library is executing the pip commend from a python console. For Unix/macOS users, use “python -m pip install ube3_apa”, and Windows users can use “python -m pip install ube3_apa” instead.

The main function that performs that analysis is e3enrich. It takes two tables and a set of parameters as standard input. The first of two tables is the site ratio table which records the protein ID, the site position, and site ratio of every ubiquitylation site in this experiment. The second one, the protein ratio table, contains information about protein ID and the ratio of different proteins instead of sites. The input of this table is optional and triggers normalization of site ratio by corresponding protein ratio. Other parameters can be modified to fit different types of protein ID inputs, change the directory that results are generated and select various output formats based on research focus. A detailed explanation of this function is in the Supplementary Information. And contents of input and output table were also explained (**Table S1**).

All related files including the code, the E3 ligase substrate dictionary, example input files were also uploaded to GitHub, which can be downloaded from the following link: https://github.com/Chenlab-UMN/Ub-E3-ligase-Activity-Profiling-Analysis.

## 3 Results

### 3.1 Establish a comprehensive ESI interaction network

We collected datasets with rich information of ESIs supported by biological experiments from multiple sources, including the UbiBrowser, the Ubinet, and another published ESI database (Li *et al.*, 2017, 2021)(Chen *et al.*, 2019). Comparing the information from the three data sources showed a largely overlapping information with some differences (**Figure 2**). After removing redundancy and discrepancies, we established an integrated database for human ubiquitin E3 ligase-substrate interaction that include 354 E3 ligases and 2501 interactions (**Table S2**). All E3 ligases interactions collected in the database were built in the package to allow comprehensive analysis.

**Figure 2.**
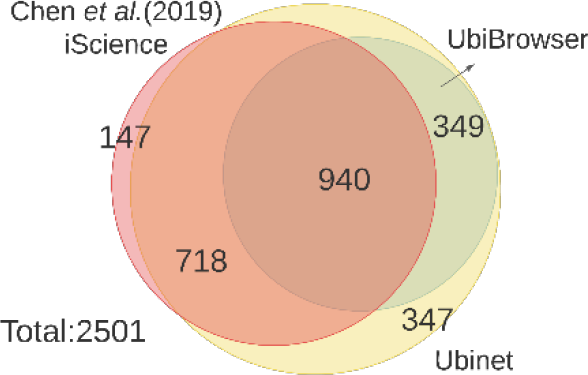
Overlap and integration of the E3 ligase-substrate data resources.

### 3.2 Profile E3 ligase activity in quantitative ubiquitylome studies

With the establishment of a comprehensive E3 ligase-substrate database, the program collects the site and protein-specific data from quantitative ubiquitylome studies. To profile E3 ligase activity based on this data, we reasoned that a physiologically meaningful changes in ubiquitin E3 ligase activities should be reflected on the overall changes of the ubiquitination abundance of their corresponding targets (**Figure 3a**). As the ubiquitiylome proteomics analysis mainly provide site-specific quantification of ubiquitination, the program offers the option to calculate the averaged site ratios of a target protein to represent the ubiquitination changes of each protein for the protein-level E3 ligase activity profile analysis. Then, all quantified ubiquitination proteins for a specific E3 ligase in the dataset will be collected as a sample group. The same number of quantified ubiquitination proteins as the number of quantified substrates for any given E3 ligase will be randomly selected from the ubiquitylome dataset in a bootstrapping procedure. Based on the Central Limit Theorem, the average quantification ratios of each group of randomly selected ubiquitinated proteins should form a normal distribution. Based on this reference distribution, we can estimate the statistical significance of the average ubiquitination ratio of the substrate group for an E3 ligase (**Figure 3a**). A significant change in the activity profile can indicate a significant increase or decrease of E3 ligase activity in the context of experimental conditions comparing to the overall changes in ubiquitination dynamics in the background. The protein-level E3 ligase activity profile analysis was used for the downstream applications.

**Figure 3.**
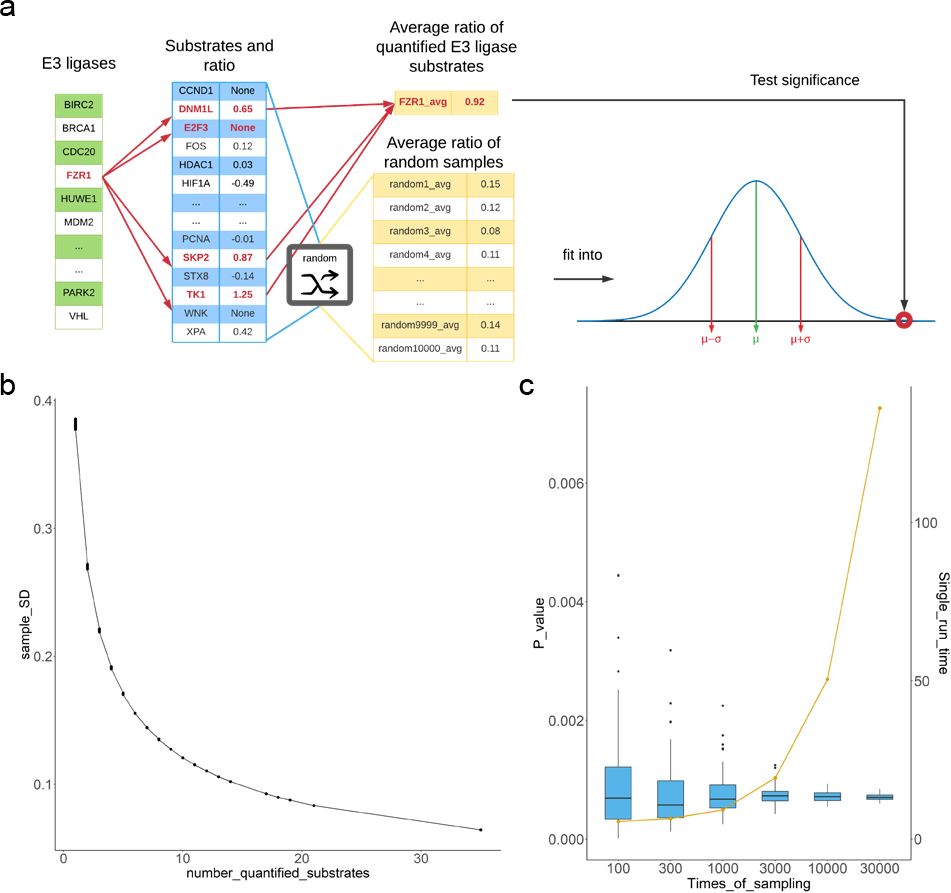
Establishing the statistical model for UbE3-APA. a. Statistical modeling to evaluate an E3 ligase activity profile in UbE3-APA analysis. b. Variations of standard deviations of the sampling ratios towards E3 ligases with different number of quantified substrates (testing with a dataset in the SPOP study). c. Variations of p-value and analysis time for single run with various number of sampling times for SPOP E3 ligase activity profiling in a dataset of the SPOP study.

To further explore how the number of quantified substrates and the times of sampling may affect the analysis processes and results, we performed tests with different parameters (**Figure 3b-c**). Our data showed that if more substrates of any given E3 ligases were quantified, the standard deviation of randomly selected sample ratios for the activity profile analysis decreased, which therefore led to more statistically significant estimation of E3 ligase substrate ratios based on the distribution (**Figure 3b**). This built-in mechanism of our model certainly supports the notion that E3 ligase activity profile could be better assessed if more E3 ligase substrates were quantified. Our test of the sampling process showed that increasing the number of random samplings in the bootstrapping process could reduce the variation of p-values calculated based on the sampling distribution and therefore led to more reliable and precise estimation of the statistical significance (**Figure 3c**). On the other hand, increasing the number of random samplings would also cost more time per run and reduce the efficiency of the analysis (Figure 3c). Considering both efficiency and reliability of the results, we have selected 10,000 times of repeats for the random selection process in E3 ligase activity profiling analysis.

### 3.3 Validate UbE3-APA workflow with the quantitative ubiquitylome analysis of SPOP E3 ligase

To validate our algorithm, we chose a quantitative proteomics study that aimed to characterize ubiquitination dynamics that was mediated by SPOP, an E3 ligase that is frequently mutated in prostate cancer and affects the regulation of downstream pathways in cancer progression (Theurillat *et al.*, 2014). This study included two sets of quantitative proteomics experiments and each set of experiment aimed to quantify the ubiquitination dynamics upon the overexpression of vector control, SPOP-wild-type (WT) and one of the two SPOP-mutants F133L and Y87N. Both mutations are naturally occurring mutations in prostate cancer and known to suppress the SPOP-WT induced ubiquitylation. The quantitative analysis was performed using SILAC workflow with the expression of each form of SPOP-WT or vector control pairing to one of the SPOP-mutant.

Using UbE3-APA workflow, we analyzed the normalized quantitative ubiquitination ratios included in their supplemental information across all six pairs of SILAC experiments (**Table S3**). The activity profiling analysis showed that the SPOP activity was significantly enriched in cells overexpressing SPOP-WT when comparing to cells overexpressing vector control or either one of the SPOP mutants (**Figure 4a**). When comparing between cells expressing SPOP mutant and vector control, our model found no significant changes in SPOP activity. These profiling analysis results matched well with SPOP ubiquitylation activity differences expected in the original study. The activity profiling analysis also allowed us to generate volcano plots with the statistical significance test and quantification ratios. Two examples with the Y87N (L)-WT (H) group and the Y87N (L) – mutant (H) group were shown (**Figure 4b-c**). As clearly shown, when SPOP-WT was overexpressed, the SPOP activity increased significantly comparing to the overexpression of SPOP-Y87N mutant with significantly increased SILAC H/L ratios of SPOP target proteins, while the SPOP activity did not change when comparing the cells overexpressing SPOP-Y87N mutant and vector control.

**Figure 4.**
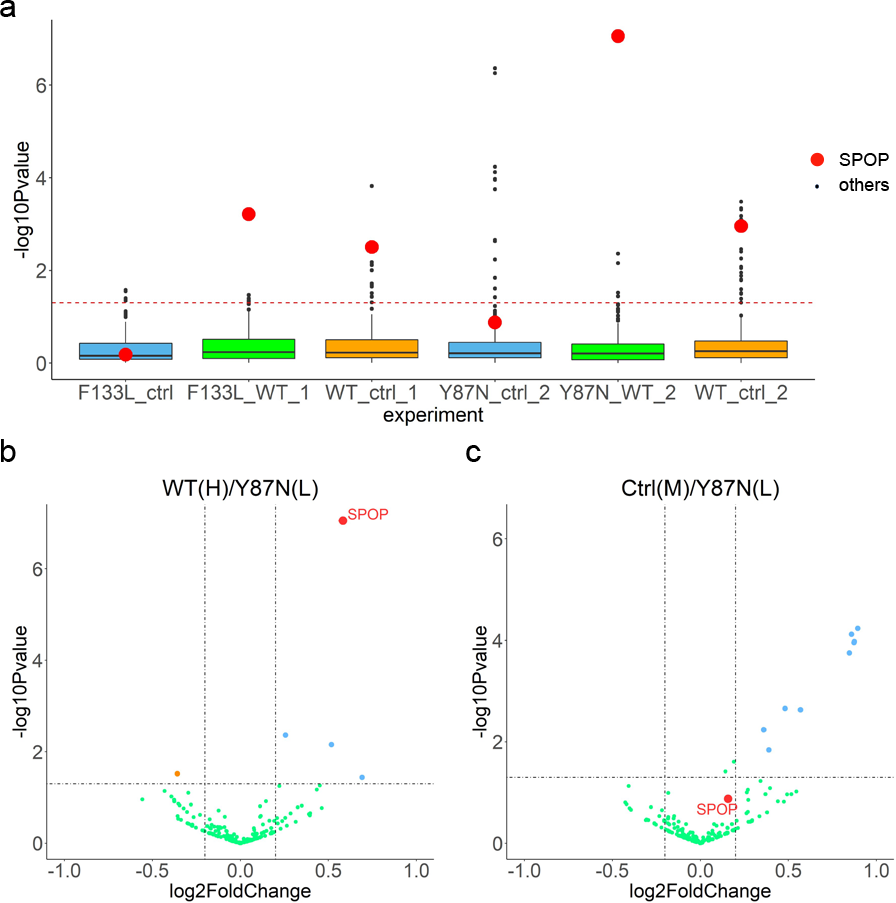
Protein level E3 ligase activity profiling results of the SPOP study. Ctrl, vector control; WT, wildtype-SPOP; 133L, SPOP-F113L; 87N, SPOP-Y87N; 1 and 2, experiment set one and two. a) All experiments of the SPOP study in box plot, each box contained enrichment p-values of all E3 ligases in one SILAC experiment. b) SPOP-Y87N to SPOP-WT group in volcano plot. c) SPOP-Y87N to vector control group in volcano plot.

### 3.4 Apply UbE3-APA model to profile E3 ligase activities in response to mitochondrial depolarization

We applied UbE3-APA workflow to analyze a quantitative ubiquitylome study that focused on PARKIN and global ubiquitylation network in response to mitochondrial depolarization (Sarraf *et al.*, 2013). This large-scale study included 73 quantitative ubquitylome proteomics analysis to explore the dynamics of the ubiquitination pathways under various mitochondrial depolarization treatment as well as in cells with different genetic background, and detailed information of experiment condition of each group we collected from supplemental data in original paper (Sarraf *et al.*, 2013) were listed (**Table S4**). Analysis of all the datasets with UbE3-APA workflow showed that when mitochondria was not damaged, there was not an apparent PARKIN activity even when PARKIN was overexpressed (**Table S5**). Once the mitochondria were polarized, there was a significant increase of PARKIN activity (**Figure 5a**). Inhibition of Pink1, the upstream kinase activating PARKIN, abolished the activation of PARKIN as expected when mitochondria was depolarized. When mitochondria were depolarized, we could not see an apparent activation of PARKIN based on ubiquitylome analysis data (**Figure 5b**). But when cells were treated with baflomycin, an autophagy inhibitor, there was a strong indication of activation of PARKIN upon mitochondria depolarization (**Figure 5c**), suggesting that the ubiquitinated substrates of PARKIN could not be efficiently degraded. Therefore, this data agrees well with the knowledge that PARKIN activation led to mitochondria degradation through autophagy process and it also suggested that PARKIN substrates are mainly degraded through autophagy pathways. In addition, the dataset also included the proteasome inhibition experiment upon mitochondria depolarization. Interestingly, our analysis showed that the inhibition of proteasome activity alone showed no strong enrichment of PARKIN substrate ubiquitination, which could suggest that either PARKIN substrates were mainly degraded through processes other than proteasome degradation (such as autophagy), or PARKIN was not activated upon mitochondria depolarization when proteasome was inhibited (**Figure 5c**). Analysis of the dataset with the cotreatment of cells with both proteasome inhibitor and autophagy inhibitor showed that PARKIN was indeed not activated upon proteasome inhibition because even autophagy inhibitor treatment failed to enrich PARKIN ubiquitination substrates (**Figure 5c**). This finding agreed well with previously published observation that upon proteasome inhibition, mitochondria depolarization failed to induce mitochondria fragmentation despite of PARKIN translocation to mitochondria (Tanaka *et al.*, 2010).

**Figure 5.**
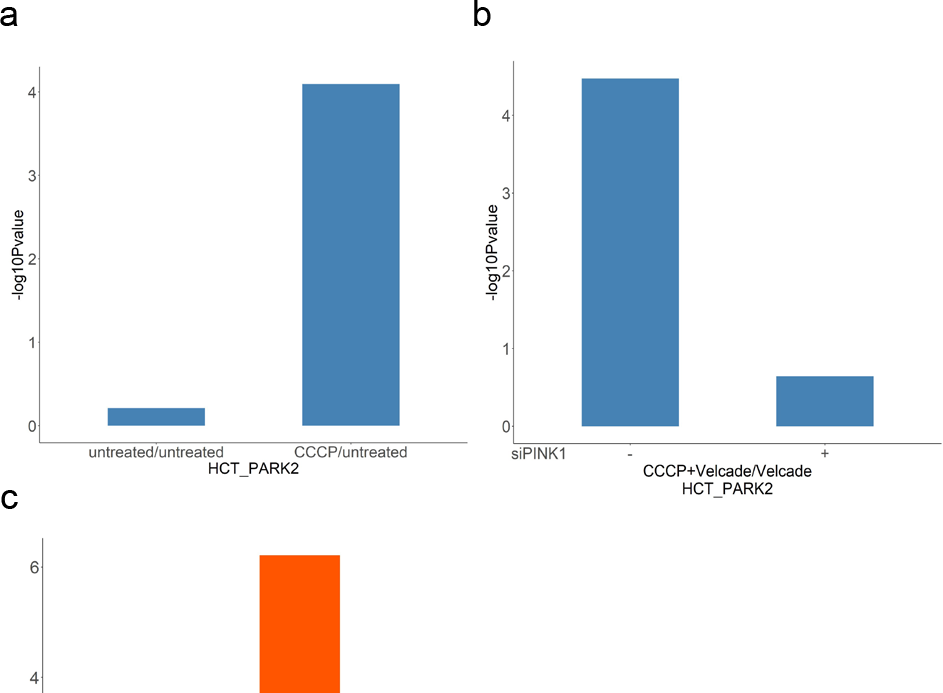
Ubiquitin E3 ligase activity profiling analysis of <u>PARKIN</u> under various genetic and chemical treatment. a. PARKIN activity difference in HCT116 cells with PARK2 overexpression and with or without carbonyl cyanide m-chlorophenyl hydrazone (CCCP, mitochondrial depolarization inducer) treatment. b. PARKIN activity difference in HCT116 cells with PARK2 overexpression and with or without Pink1 inhibition. c. PARKIN activity difference in HCT116 cells with the treatment of bafolmycin (BafA, an autophagy inhibitor) and/or Velcade (a proteo-some inhibitor).

We then integrated the analysis of the ubiquitylome dynamics in all experimental conditions in the mitochondria depolarization study and plot the E3 ligase activity profiles in heatmap (**Figure 6, Table S6**). The E3 ligases were clustered with hierarchical clustering based on how similar their activity profiles change under different treatment conditions. Out of 203 E3 ligases profiled by our model across all experimental conditions, three E3 ligases (FZR1, AMFR, MARCHF5) showed a very similar activity pattern as PARKIN across most of the experimental conditions. Since the activity profiles of E3 ligases were analyzed based on their corresponding substrates, it was likely that E3 ligases showed similar activity profiles when they shared common substrates. For better clarification, we mapped the E3-ligase - substrate interaction networks for the four E3 ligases (**Figure 7a, Table S7**). The network indicated the shared and unique connection between each E3 ligase and corresponding substrates, the number of times the substrates were quantified under all conditions and the significance of ubiquitination level changes for the substrate proteins. We can clearly see that the unique substrates of AMFR and FZR1 did not change significantly to contribute to activity profiles and their activity changes were mainly caused by the changes of ubiquitination levels in substrates shared with PARKIN. Only MARCHF5 had unique substrates whose ratios were changed significantly and similarly along with those unique substrates of PARKIN. For better clarification of the data, we included “Group” mode to the result output. In this mode, the software will group E3 ligases that share substrates together if the E3 ligases do not have unique substrate and only the E3 ligases that contain all the substrates in the group were labeled as leading E3 ligases of the group. We re-analyzed all the data using the Group mode and identified 127 E3 ligase groups (**Table S8**). Then, we performed correlation analysis of all the E3 ligase groups in distance matrix. The data clearly showed that MARCHF5 and PARKIN shared the most similar activation profiles (**Figure 7b**). This finding confirmed that MARCHF5 was also activated during mitochondrial depolarization, which agrees well with the previous finding that PARKIN-dependent ubiquitination targets MARCHF5 for translocation and activation (Koyano *et al.*, 2019).

**Figure 6.**
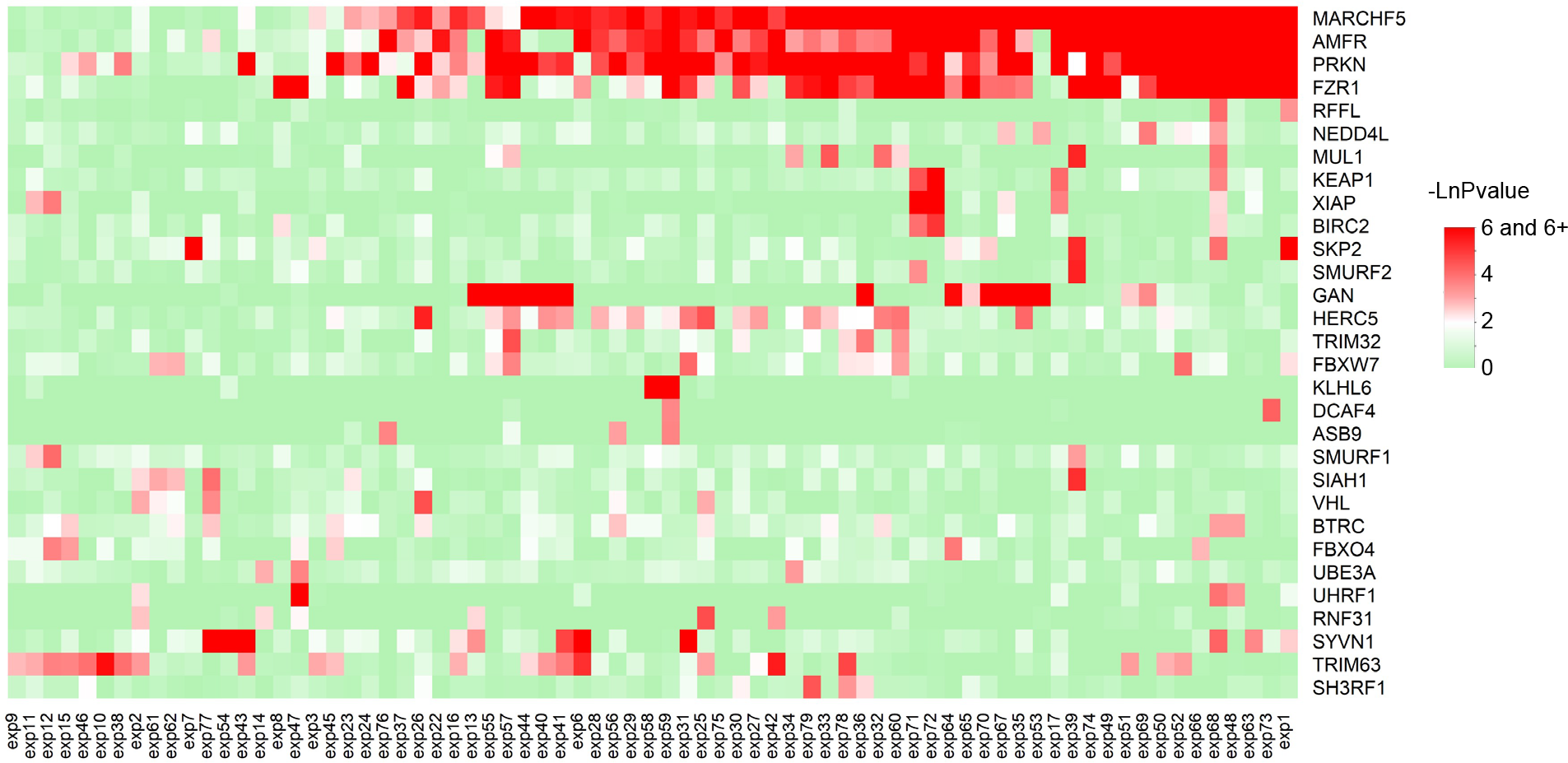
E3 ligase co-activation profiles across all experiments involved in the mitochondria depolarization study. The p value enrichment of <u>E3 ligases across 73 groups of experiments were -ln transformed. Only E3 ligases that were enriched (p<0.05) in at least one experiment were included in this heatmap</u>.

**Figure 7.**
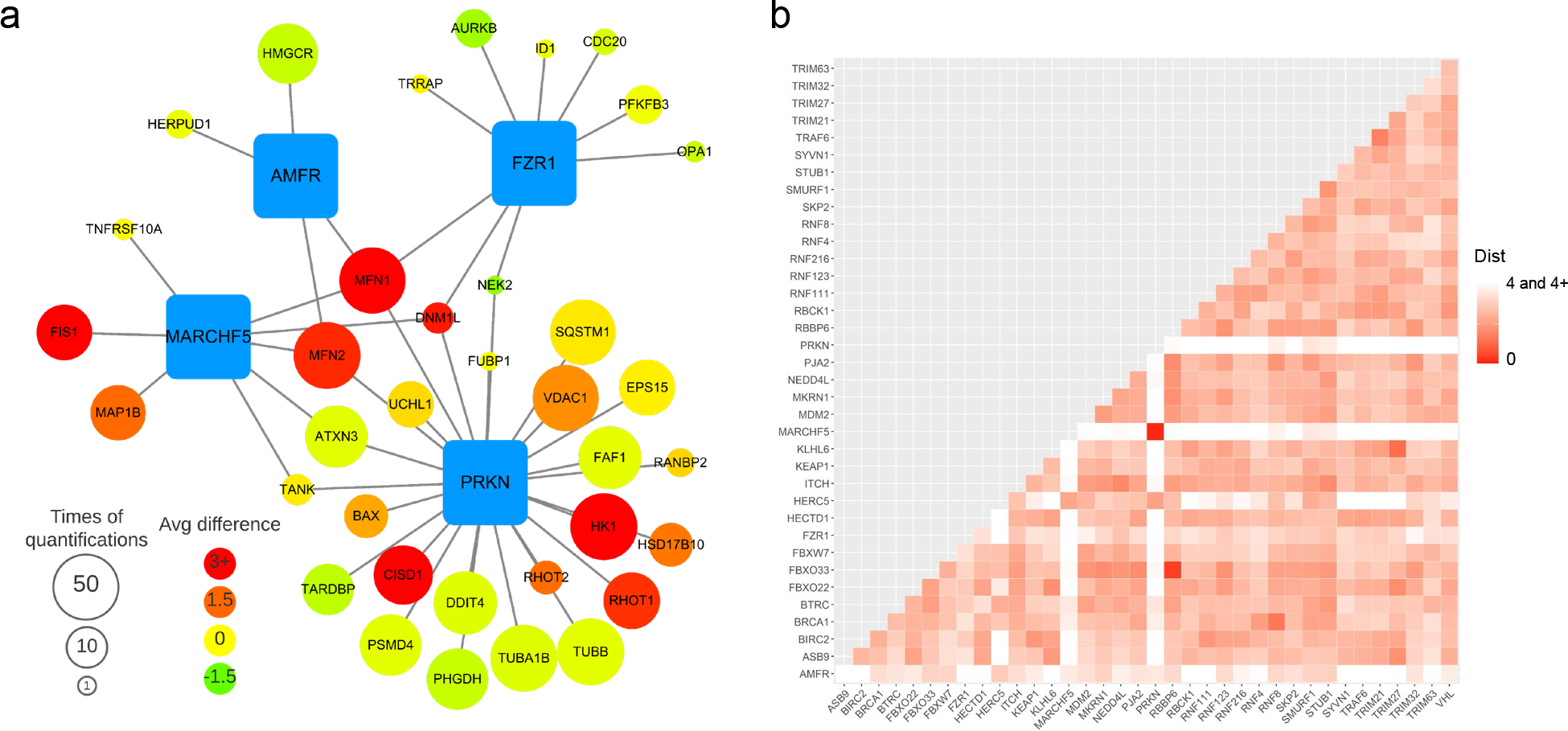
Regulation <u>networks</u> between four E3 ligase, PARKIN, MARCHF5, AMFR and FZR1 in the PARKIN study. a) Interaction network of four E3 ligases and their substrates found in the mitochondria depolarization experiments. Blue square represents E3 ligases, circle represents substrates, edge represents Enzyme-substrate Interaction (ESI), size of circles indicates <u>the number of groups (out of 73 experimental groups) this substrate being quantified and used for the corresponding E3 ligase activity analysis, and the color of the circle indicates the difference in average log2 quantification ratio between the substrate ratio and the group average</u>. b) <u>Correlation matrix and heatmap</u> of leading E3 ligase activity profiles from all experiments. <u>Color gradient represents 2-D Euclidean distance (Dist) between a pair of leading E3 ligase activity profiles. Only E3 ligases with p < 0.2 in at least one experiment were included</u>.

### 3.5 Apply UbE3-APA model to Data Independent Acquisition (DIA) Dataset

To further explore the usage of our UbE3-APA model, we applied it to two recently published quantitative ubiquitylome studies based on DIA analysis. The first study explored how the ubiquiyltome was associated with the TNF signaling pathways (Hansen *et al.*, 2021b) by treating cells with or without TNF. We collected site-specific intensities of all replicates in treated group and mock group and calculated the intensity ratios between TNF treated and mock treated cells. Then we performed UbE3-APA analysis on the protein level in the grouped mode. The activity profiling analysis revealed several distinct up- and down-regulated E3 ligases under TNF treatment (**Figure 8, Table S9**). Among these up-regulated E3 ligases, TRAF2 and TRAF6 were members of tumor necrosis factor receptor-associated factors whose ubiquitination activity was crucial in the TNF signaling pathways (Bradley and Pober, 2001). In addition our analysis also identified up-regulation of activity for RNF216, SOCS3 and down-regulation of activity for MIB1 and FBXO33.

**Figure 8.**
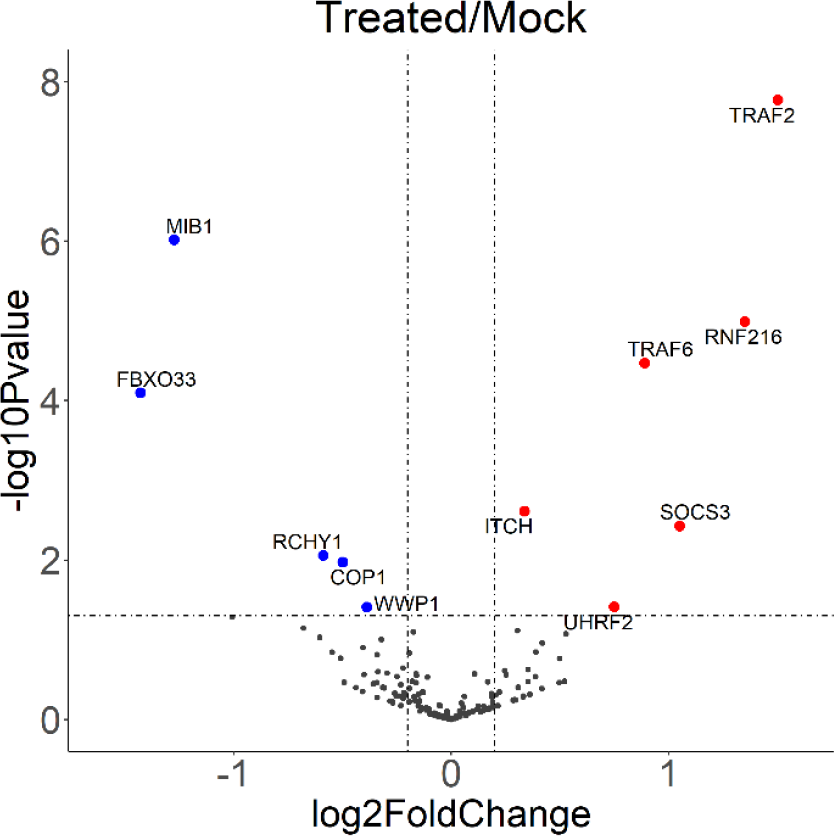
E3 ligase activation analysis based on the ubiquitination dynamics between TNF treated and mock treated cells in the TNF study.

The second study investigated the ubiquitylome changes in response to the inhibition of deubiquitinase USP7 by siRNA knockdown or chemical inhibitor such as FT671 (Steger *et al.*, 2021). We extracted their DIA-based ubiquitylome data under these four conditions: siCTRL+DMSO, siCTRL+FT671, siUSP7+DMSO, and siUSP7+FT671 and then calculated the ratio difference between FT671 and DMSO treatments under either siCtrl background or siUSP7 background. Analyzing the two pairs of ubiquitylome datasets with UbE3-APA revealed differential activation profiles of E3 ligases upon the FT671 treatment with siUSP7 background or siCTRL background (**Figure S1, Table S10**). In agreement with the findings in the published study, more E3 ligases showed altered activity profiles upon FT671 treatment but these activity changes were attenuated under siUSP7 background, suggesting that FT671 treatment was specific in targeting USP7 activity in cells.

## 4 Discussion

Advances in quantitative proteomics have enabled large-scale profiling of ubiquitination substrates, also known as the ubiquitylome. Application of quantitative proteomics in ubiquitylome analysis revealed the key ubiquitination targets in the biological processes and determined the down-stream signaling pathways that were most significantly affected by the ubiquitination process. Yet, few studies systematically examine the upstream regulatory pathways of the ubiquitination. Analysis of regulatory enzyme activities has been largely limited to a few well-selected targets of each enzyme. Recent advances in the collection and biological validation of E3 ligase-substrate database provide a great opportunity to use ubiquitylome quantitative analysis as an activity-readout to profile the ubiquitin E3 ligases.

In this study, we developed a statistical framework and workflow to identify the ubiquitin E3 ligase activity in a high-throughput and unbiased manner. This open-source python package enabled effective profiling of E3 ligase activities through robust statistical analysis based on quantitative ubiquitylation results. In the case study of SPOP E3 ligase ubquitylome analysis, our model correctly validated the SPOP activity upon the over-expression of SPOP WT and mutant forms with various activity. Application of our workflow to analyze the ubiquitylome dynamics upon mitochondria depolarization confirmed that activation of PARKIN E3 ligases under various conditions and unexpectedly discovered the role of proteasome inhibition on PARKIN activation. Our statistical framework allowed us to collect the E3 ligase activity profiles across multiple conditions. Application of our workflow to profile 73 quantitative ubiquitylome analysis enabled clustering analysis of E3 ligases across the experimental conditions and revealed the co-activation of PARKIN and MARCHF5 upon mitochondrial depolarization. Our mothods was further applied to two studies with DIA ubiquitomes, and in both case studies, E3 ligases related with the treatment proved by previous papers were revealed by our model through activation profile changes.

The statistical framework described in this study can be generally applied to other PTM pathway analysis. We have recently applied the workflow and developed Kinase Activity Profiling Analysis (KAPA) to identify iron deficiency induced activation of AMPK pathway in neuronal cells (Erber *et al.*, 2021). Our study demonstrated that it is possible to apply statistical analysis workflow to systematically profile E3 ligase activity.

However, we also recognize that efficient analysis of E3 ligase activity in a system-wide manner is limited by several factors. First, the number of E3 ligase-substrate interaction in our knowledgebase is still limited comparing to other PTMs such as phosphorylation and acetylation. Our integrated database from various sources contained 2354 gene-level interactions of humans in total, which is quite small comparing to 13855 gene-level interactions between phosphorylation sites and kinases collected by PhosphoSitePlus in human (Hornbeck *et al.*, 2015). Continued effort in the high throughput discovery of E3 ligase and substrate interaction is needed to expand the knowledgebase for more reliable and confident analysis of upstream regulatory enzyme activities.

Secondly, although current high throughput proteomics have allowed in-depth quantification of ubiquitylome in single experiment, the data-dependent analysis (DDA) often suffers from limited reproducibility and reduced quantification precision. For example, in our analysis of mitochondria depolarization ubiquitylome study, for E3 ligase FZR1 and AMFR, they have 56 and 14 substrate proteins respectively based on our ESI database, but only 10 and 4 substrates were found at least once in all 73 groups of experiments. Therefore, their activity profiles were affected by the shared substrates with PARKIN. If more substrates were reproducibly quantified, the analysis profiles of the two E3 ligases could be more accurate and informative. Application of data independent acquisition (DIA) for ubiquitylome analysis as we demonstrated in our study will certainly help address this challenge (Hansen *et al.*, 2021a).

Lastly, currently E3 ligase and substrate interaction database has been largely based on the protein-level and there is limited knowledge on the site-specificity of ubiquitination regulatory pathways comparing to the knowledge on the kinase-phosphorylation regulatory network. Lack of site-specific regulation information presents a challenge to reveal potential regulatory enzyme activities on overlapping protein substrates but on distinct target sites., making it less precise compared to other well-studied PTMs based on the regulatory network between enzymes and sites, for example, phosphorylation and acetylation. It requires the continued development technologies to identify major enzyme target sites in the ubiquitination pathway. Future updates of the program will include updated E3 ligase-substrate interactions database with the potential for site-specific enzyme activity analysis.

## Acknowledgements

We would like to thank Gaurav Behera and other members of the Chen lab for helpful suggestions and discussions.

