## Supplemental Information for "UbE3-APA: A Bioinformatic Strategy to Elucidate Ubiquitin E3 Ligase Activities in Quantitative Proteomics Study"

#### Instructions for software installation and usage

#### Supplemental Figure

**Figure S1.** E3 ligase activity profiling results of the USP7 study. a) Volcano plot of E3 ligase activity profiles comparing FT671+siCTRL and DMSO+siCTRL treatment conditions and b) volcano plot of E3 ligase activity profiles comparing FT671+siUSP7 and DMSO+siUSP7.

#### Supplemental Tables

**Table S1.** Contents of Input and Output table columns of Ub E3 ligase Activity Profiling Analysis.

**Table S2.** Human E3 ligase interaction database integrated from three sources.

**Table S3.** Ub E3 ligase Activity Profiling Analysis of the SPOP mutant and wildtype ubiquitylome study.

**Table S4.** Descriptions of experimental condition for mitochondrial depolarization and PARKIN activity studies

**Table S5.** Ub E3 ligase Activity Profiling Analysis of the quantitative ubiquitylome analysis of mitochondrial depolarization and PARKIN activity.

**Table S6.** Enrichment matrix of E3 ligases from Ub E3 ligase Activity Profiling Analysis of mitochondrial depolarization and PARKIN activity

**Table S7.** Integrated properties and interaction network of four E3 ligases and their substrates from Ub E3 ligase Activity Profiling Analysis of mitochondrial depolarization and PARKIN activity in grouped mode

**Table S8.** Ub E3 ligase Activity Profiling Analysis of the quantitative ubiquitylome analysis of mitochondrial depolarization and PARKIN activity in grouped mode.

**Table S9.** Ub E3 ligase Activity Profiling Analysis of the quantitative ubiquitylome analysis of TNF treatment study in grouped mode.

**Table S10.** Ub E3 ligase Activity Profiling Analysis of the quantitative ubiquitylome analysis of USP7 inhibition study in grouped mode.

### Instructions for software installation and usage

Our ube3_apa python package can be installed through executing the following command in a python console:

```
python3 -m pip install ube3_apa
```

After successful installation, you may test the code with the testing data on our GitHub website: https://github.com/Chenlab-UMN/Ub-E3-ligase-Activity-Profiling-Analysis

There are some example codes:

```
#########################################

import ube3_apa

input_directory = “directory_of_testdata_folder”

#standard

ube3_apa.e3enrich(siteratio_dir=input_directory+”/testdata/siteratio_testdata.csv”, input_type=”UniprotAC”, output_dir=”desired_output_directory”, exp_label=”1”, grouped=False, output_ratio=True, log2trans=False)

#grouped

ube3_apa.e3enrich(siteratio_dir=input_directory+”/testdata/siteratio_testdata.csv”, input_type=”UniprotAC”, output_dir=”desired_output_directory”, exp_label=”2”, grouped=True, output_ratio=True, log2trans=False)


#with protein normalization

ube3_apa.e3enrich(siteratio_dir=input_directory+”/testdata/siteratio_testdata.csv”, input_type=”UniprotAC”, output_dir=”desired_output_directory”, exp_label=”3”, proratio_dir=input_directory+”/testdata/proteinratio_testdata.csv”, grouped=False, output_ratio=True, log2trans=False)

#########################################
```

After running the code above, you will find some csv files in the output directory. The enrichment p-values are listed in files initiated with “UbE3_APA”, and there are also other related data such as the number of substrates found and average site ratios included.

#### Parameters of e3enrich

*e3enrich(siteratio_dir, input_type, output_dir, exp_label='', proratio_dir="None", grouped=False, output_ratio=False, log2trans=True*)

Perform Ub E3 ligase activity profiling analysis based on ratio of E3 ligase substrates

Parameters can be customized through key-value pairs as below.

Parameters:

*siteratio_dir: string*

The directory of the file that contains the ratio of every site.

*input_type: {“UniprotAC” or “protein”, “gene symbol” or “gene”}*

The type of IDs used in the site ratio file and the protein ratio file, format in both files should be the same.

"UniprotAC" or "protein"   example: Q00987, P40337, Q9HAU4, O43791

"gene symbol" or "gene"   example: MDM2, VHL, SMUF2, SPOP

List of valid input and examples will be shown if it is an invalid value.

*output_dir: string*

The directory where E3 enrichment result files will be generated.

*exp_label: string, default ””*

The string attached to the output file name that separates different results when there are multiple groups.

*proratio_dir: string, default ””*

The directory of the file that contains the ratio of every protein. A valid directory input here will trigger normalization by corresponding protein ratio for all files in this run. By default, the output will not be normalized by protein ratio.

*grouped: bool, default False*

If True, enrichment results will show leading E3 ligase and grouped E3 ligase in each row of the result instead of showing each E3 ligase individually. E3 ligases are grouped according to the relationship of the detected substrate in this run.

*ratio_output: bool, default False*

If True, files that contain the ratio of each ubiquitylation site and the average ratio of each ubiquitinated protein will be generated.

*log2trans: bool, default True*

If True, ratios will take transform y=log2(x) before the E3 ligase enrichment analysis. If False, the ratio from files will be used for enrichment analysis directly.

Returns:

This function generates files based on E3 ligase enrichment results and does not have any returns.

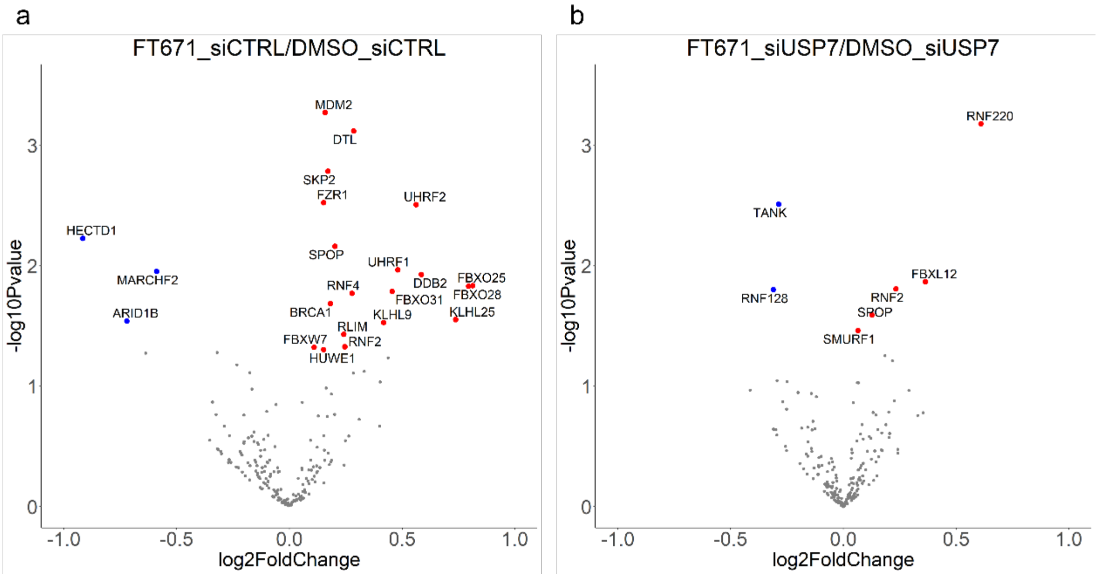


**Figure S1. E3 ligase activity profiling results of the USP7 study.** a) Volcano plot of E3 ligase activity profiles comparing FT671+siCTRL and DMSO+siCTRL treatment conditions and b) volcano plot of E3 ligase activity profiles comparing FT671+siUSP7 and DMSO+siUSP7.
